# Transient telomerase inhibition alters cell cycle kinetics

**DOI:** 10.1101/158287

**Authors:** Connor AH Thompson, Alice Gu, Sunny Yang, Veena Mathew, Helen B Fleisig, Judy MY Wong

## Abstract

Telomerase is the ribonucleoprotein reverse transcriptase that catalyzes the synthesis of telomeres at the ends of linear human chromosomes and contributes to proper telomere-loop (T-loop) formation. Formation of the T-loop, an obligate step before cell division can proceed, requires the generation of a 3’-overhang on the G-rich strand of telomeric DNA via telomerase or C-strand specific nucleases. Here, we discover telomerase activity is critical for efficient cell cycle progression using transient chemical inhibition by the telomerase inhibitor imetelstat. Telomerase inhibition caused changes in cell cycle kinetics and increased the proportion of cells in G2 phase, suggesting delayed clearance through this checkpoint. Investigating the possible contribution of unstructured telomere ends to these cell cycle distribution changes, we observed that imetelstat treatment induced γH2AX DNA damage foci in a subset of telomerase-positive cells but not telomerase-negative primary human fibroblasts. Chromatin-immunoprecipitation with γH2AX antibodies demonstrated imetelstat treatment-dependent enrichment of this DNA damage marker at telomeres. Notably, the effects of telomerase inhibition on cell cycle profile alterations were abrogated by pharmacological inhibition of the DNA-damage-repair transducer ATM.

Additionally, imetelstat potentiation of etoposide, a DNA-damaging drug that acts preferentially during S/G2 phases of the cell cycle, also depended on functional ATM signaling. Our results suggest that telomerase inhibition delays the removal of ATM-dependent DNA damage signals from telomeres in telomerase-positive cancer cells. This demonstrates for the first time that telomerase activity directly facilitates the progression of the cell cycle through modulation of transient telomere dysfunction signals.

## INTRODUCTION

Telomeres are nucleoprotein structures present at the ends of eukaryotic linear chromosomes. Telomeres differentiate the ends of chromosomes from random DNA breaks through the formation of capping structures called telomere loops (T-loops). These higher-order chromatin structures are essential for proper telomere function and cell cycle progression. In mammals, telomeres are composed of repeated sequences of (TTAGGG)_n_ nucleotides, the complementary DNA strands, and associated proteins (1). Telomeres prevent the loss of coding DNA sequence by buffering the lagging strand gap left by the removal of RNA primers, thereby solving the “end replication problem”. Telomeric DNA loss also results from T-loop resolution, an obligatory step to provide access for DNA replication machinery. Together, these processes contribute to the loss of 50-100 base pairs of 3’ terminal telomeric DNA with each replication cycle (2). This telomere attrition forms the basis of the Hayflick Limit and restricts the number of times a cell lineage may proliferate (3).

Telomeres are maintained by the ribonucleoprotein telomerase. Telomerase has two main components: the catalytic telomerase reverse transcriptase (TERT) and the template telomerase RNA (TER). TERT catalyzes the synthesis of the hexanucleotide repeats by reverse transcribing the RNA template sequences encoded in TER (4-7). This telomerase-mediated telomere elongation promotes immortal growth by decoupling the cell division limit from telomere length attrition.

TERT expression is normally repressed or only transiently activated in somatic human cells, but telomerase activation and constitutive expression is detected in 85-90% of human cancers (8,9). These differences make telomerase inhibition an attractive therapeutic target (1,5-7,10,11,12). One current strategy for telomerase inhibition is the drug imetelstat (GRN163L), a synthetic, lipid-conjugated, 13-mer oligonucleotide N3’ P5’ thiophosphoramidate complementary to the template of the TER component of telomerase (4,11,13,14-17). As a competitive antagonist, imetelstat blocks the normal association of TER with chromosomal ends (substrates), efficiently inhibiting *de novo* telomeric repeat synthesis.

During the DNA-synthesis phase of the cell cycle, the activities of multiple RecQ helicases, exonucleases, homologous recombination pathway effectors, and histone methylation enzymes are coordinated to allow DNA polymerase and telomerase (when expressed) to copy through T-loops and then restore the heterochromatin state of newly replicated telomeres (8,10-11,13,18-19). Additionally, the correct rebuilding of chromosomal-end structures prior to cell division has been shown to involve the Ataxia Telangiectasia Mutated (ATM) signal transduction pathway (5-7,17,20). Activation of ATM and its associated PIKK-family member, the ATM-related (ATR) kinase, is concurrent with T-loop resolution and the resulting chromosome termini exposure during DNA replication (8,9,10-12). As a consequence, cell-cycle progression through S/G2 phase is dependent on the conclusion of proper T-loop reformation.

Previous work from our laboratory demonstrated that recombinant expression of telomerase in telomerase-negative ALT (alternative lengthening of telomeres) cells conferred a growth advantage and faster passage through S/G2 phases of the cell cycle (17,20-22). This positive role of telomerase in promoting faster cell cycle kinetics has been inferred from its ability to synthesize G-rich telomeric repeats, as expression of catalytically-dead TERT and TERT mutants that cannot recognize chromosomal end substrates was not able to support this role. However, this positive role of telomerase in promoting cell cycle progression has not been demonstrated experimentally in clinically relevant cancer models.

Here, we used imetelstat-induced chemical inhibition of telomerase activity to explore the role of *de novo* telomeric DNA repeat synthesis in cell cycle progression. Since chemical inhibition is transient, we were able to study the effects of telomerase activity on structure without significant shortening of telomere length. Transient chemical inhibition also allows the immediate observation of treatment effects, thus reducing the impact of compensatory mechanisms that may confound observations.

## RESULTS

### Telomerase inhibition increases the proportion of cells in G2/M phases of the cell cycle in telomerase-positive but not telomerase-negative cell

During normal DNA replication, DNA-damage-response (DDR) signaling is found transiently in late S/G2 phases at unstructured, open chromosome ends. These signals have been detected in the forms of ATM-MRN and ATR-ATRIP complexes (5,8,20,23-24). In order for cells to divide properly, these telomere dysfunction signals must first be cleared by the reformation of T-loops. To understand how telomerase may affect the kinetics of this process, we measured the effects of imetelstat-induced telomerase inhibition on cell cycle progression.

Cells were treated with 10 µM inhibitory doses of imetelstat (5) or its mismatch oligo control (MM) (25) for 24 h before harvesting and staining with propidium iodide (PI) for cell cycle profiling using fluorescence-activated cell sorting (FACS). In telomerase-positive mammary adenocarcinoma (MCF-7 and MDA-MB 231) and colorectal carcinoma (HT29 and LS180) cells, a significant increase in cell populations with 4N DNA content following imetelstat but not MM treatment was observed (Fig. 1 a-d). In contrast, the cell cycle profiles of telomerase-negative transformed cells (VA-13), which maintain telomeres via the alternative lengthening of telomere (ALT) mechanism (26), and primary human foreskin fibroblasts (BJ) were unaltered by both imetelstat and MM (Fig. 1 e-f).

**Figure 1.**
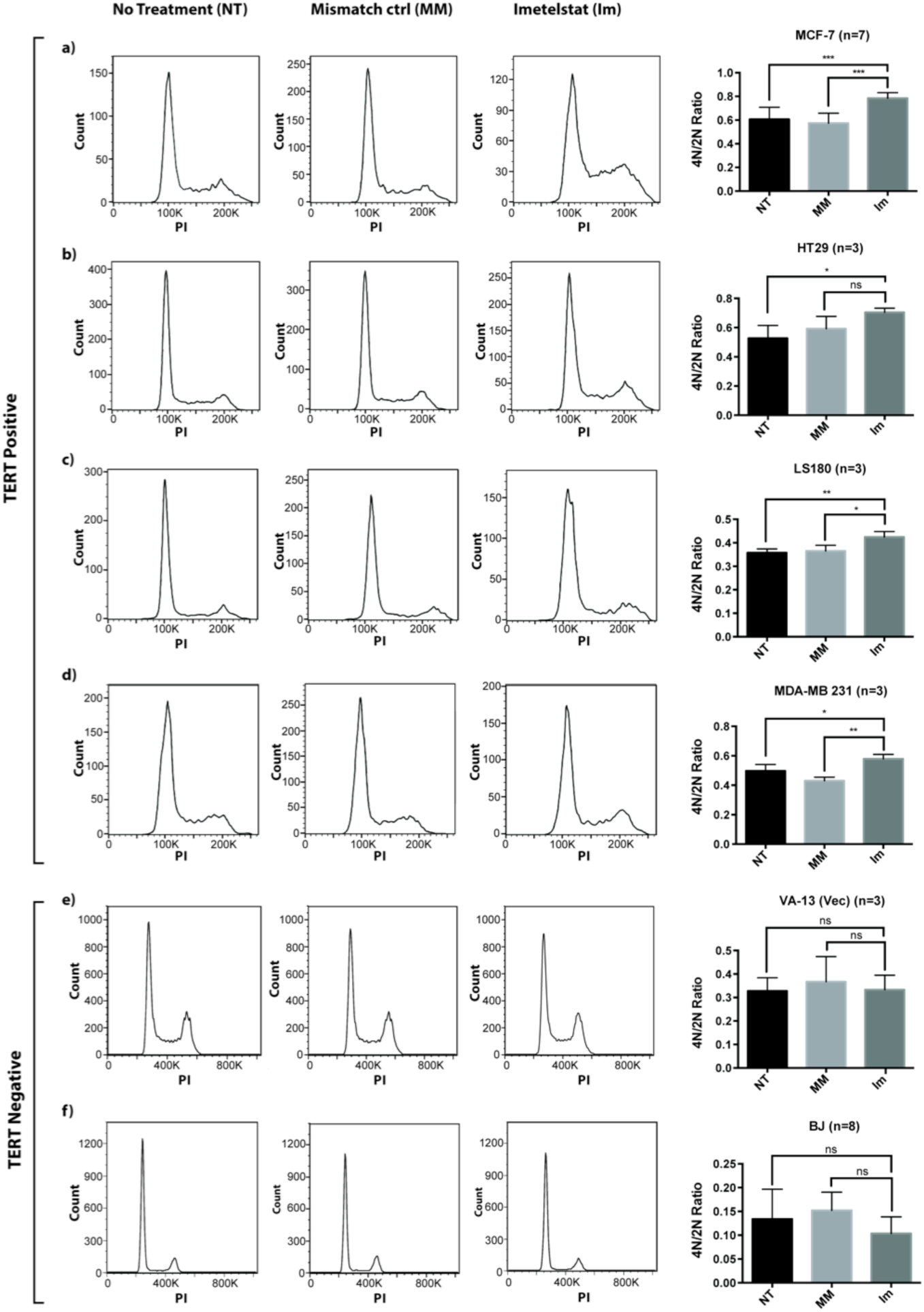
Telomerase inhibition increases the proportion of telomerase-positive cells with 4N DNA content. Cells were treated for 24 h with 10 µM imetelstat (Im) or its mismatch oligo control (MM). DNA content was analyzed using FACS and gating single cells by propidium iodide (PI) staining. Representative cell cycle profiles are shown for the indicated treatments and cell lines. (a-d) Imetelstat-induced telomerase inhibition increased the ratio of 4N to 2N cells in MCF-7, HT29, LS180, and MDA-MB 231 cell lines. (e-f) No change in ratio was observed in telomerase-negative VA-13 and BJ cell lines. ANOVA and Fisher’s LSD test were used to generate P-values (* = P ≤0.05, ** = P ≤ 0.01, *** = P ≤ 0.001). Error bars represent SD.

The increased 4N cell population was likely due to a delay in exit from G2 phase rather than a cell cycle arrest, as previous studies have reported normal cell growth following the removal of imetelstat (5). This suggests that treated cells can return to normal cycling after the removal of drugs. To confirm a delayed exit from G2 phase, we further investigated the proliferative capacity of cells after imetelstat treatment. We used the IncuCyte Zoom live cell imaging system and red fluorescent NucLight-tagged MDA-MB 231 cells to measure the effects of continuous imetelstat treatment on cell proliferation over 7 days. Consistent with a small but accumulative growth disadvantage conferred by delayed clearance from G2 phase, we observed lower nuclear counts in cells exposed to 10 µM imetelstat compared to untreated cells (Supplementary Fig. 1 a). A lower dose of imetelstat (2 µM) resulted in an intermediate growth effect that is in agreement with reduced inhibition of cellular telomerase activity (Supplementary Fig. 2, left panel).

**Figure 2.**
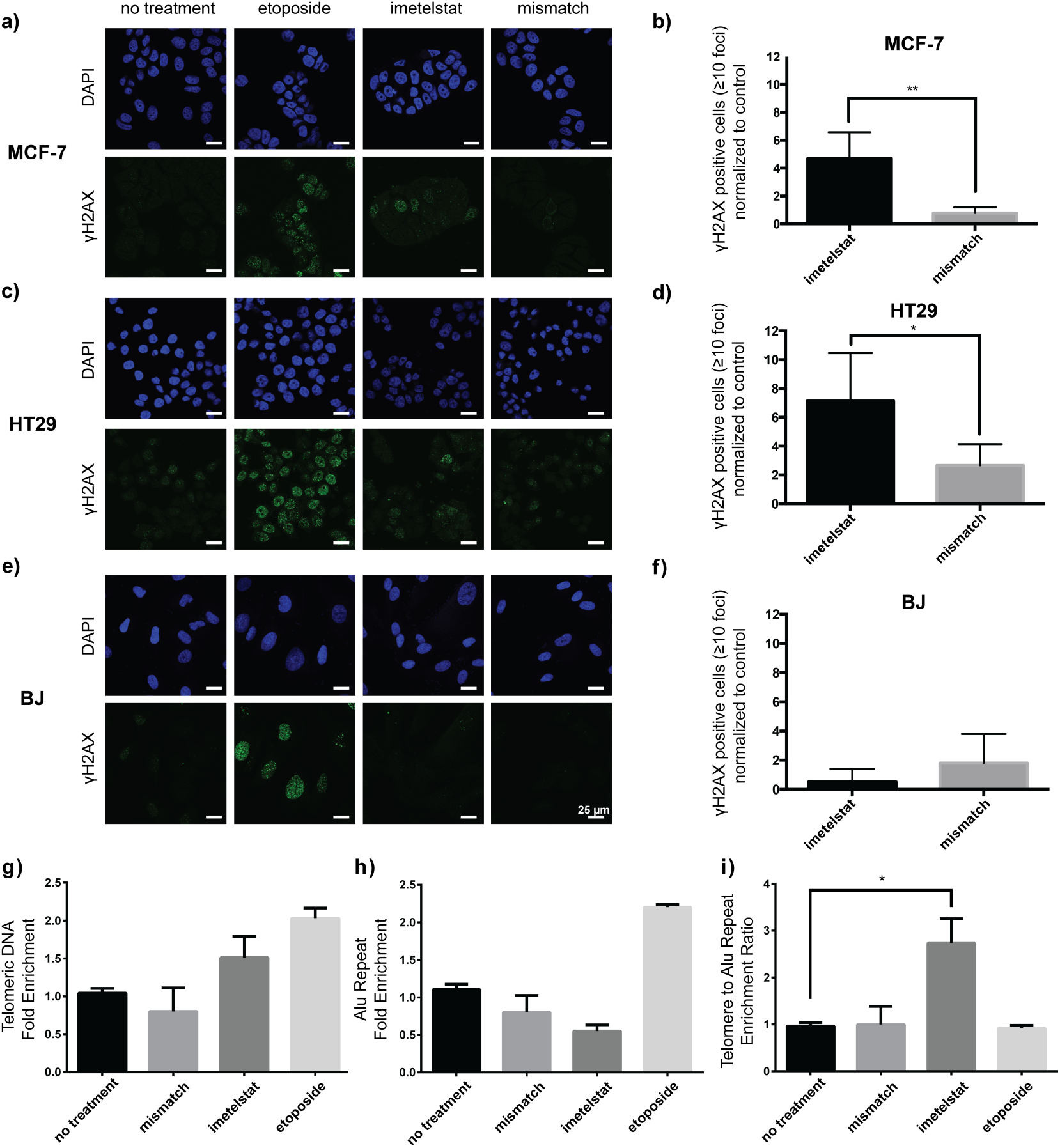
Telomerase inhibition induces DNA damage foci in telomerase-positive but not telomerase-negative cells. (a-d) Immunocytochemistry showed increased γH2AX DNA damage foci in a subset of telomerase-positive MCF-7 and HT29 cells treated for 24 h with imetelstat (10 µM). The mismatch control oligo (10 µM) did not induce this effect. Etoposide (1.4 µM) was used as a positive control. (e-f) Imetelstat did not induce DNA damage foci in telomerase-negative BJ primary human fibroblasts. Histograms show accumulation of cells with ≥10 foci, normalized to the numbers obtained from no treatment controls. Error bars represent SD. (g-i) γH2AX-ChIP-qPCR of (g) telomeric region and (h) Alu repeat region in MCF-7 cells. Enrichment values were normalized to beads only controls. (i) The ratio of ChIP enrichment of telomeric region over the Alu reference regions was indicative of telomere-specific enrichment of γH2AX signals (i). Error bars represent SEM. Student’s *t*-test was used to generate P-values (* = P ≤0.05, ** = P ≤ 0.01, n ≥ 3).

In order to measure growth in MCF-7 cells, we used the same treatment regimen and measured absolute cell counts by Coulter counting. Similar to before, a subtle growth defect was observed in MCF-7 cells treated with 10 µM imetelstat (Supplementary Fig. 1 b). In contrast, imetelstat had no apparent effect on cellular proliferation of telomerase-negative VA-13 cells (Supplementary Fig. 1 c). Telomerase activity in MDA-MB 231, MCF-7, and HT29 cells was effectively inhibited by imetelstat in a dose-dependent manner, as confirmed by telomeric repeat amplification protocol (TRAP) assays (Supplementary Fig. 2). These results are consistent with a subtle imetelstat-induced defect in cell growth seen under longer-term incubation conditions. Together, our data suggest that transient telomerase inhibition causes changes in cell cycle kinetics that result in a growth disadvantage.

### Imetelstat treatment induces telomere-specific DNA damage foci during the G2 phase of telomerase-positive cells

Imetelstat treatment altered cell cycle profiles and delayed the passage of telomerase-positive cells through G2 phase. We reasoned that this observation is linked to an imetelstat-induced delay in proper telomere structure reformation. To confirm telomerase inhibition affects the normal kinetics of telomere capping, we quantified the accumulations of telomere-specific DNA damage signals following imetelstat treatment. As uncapped telomeres are recognized as DNA damage in the absence of proper T-loop formation, we used immunocytochemistry and confocal microscopy to visualize γH2AX (Ser139), a standard DDR marker of uncapped or dysfunctional telomeres (1,17,20,23-24).

Imetelstat treatment over 24 h increased the number of cells with γH2AX DDR foci in a subset of telomerase-positive MCF-7 and HT29 cells (Fig. 2 a-d) but not telomerase-negative primary human BJ fibroblasts (Fig. 2 e-f). In parallel, the mismatch oligo control had no significant effect on γH2AX DDR foci accumulation. The observed increase in foci was not due to an increase in non-specific background labeling in imetelstat-treated cells as illustrated by the secondary antibody-alone controls (Supplementary Fig. 3).

**Figure 3.**
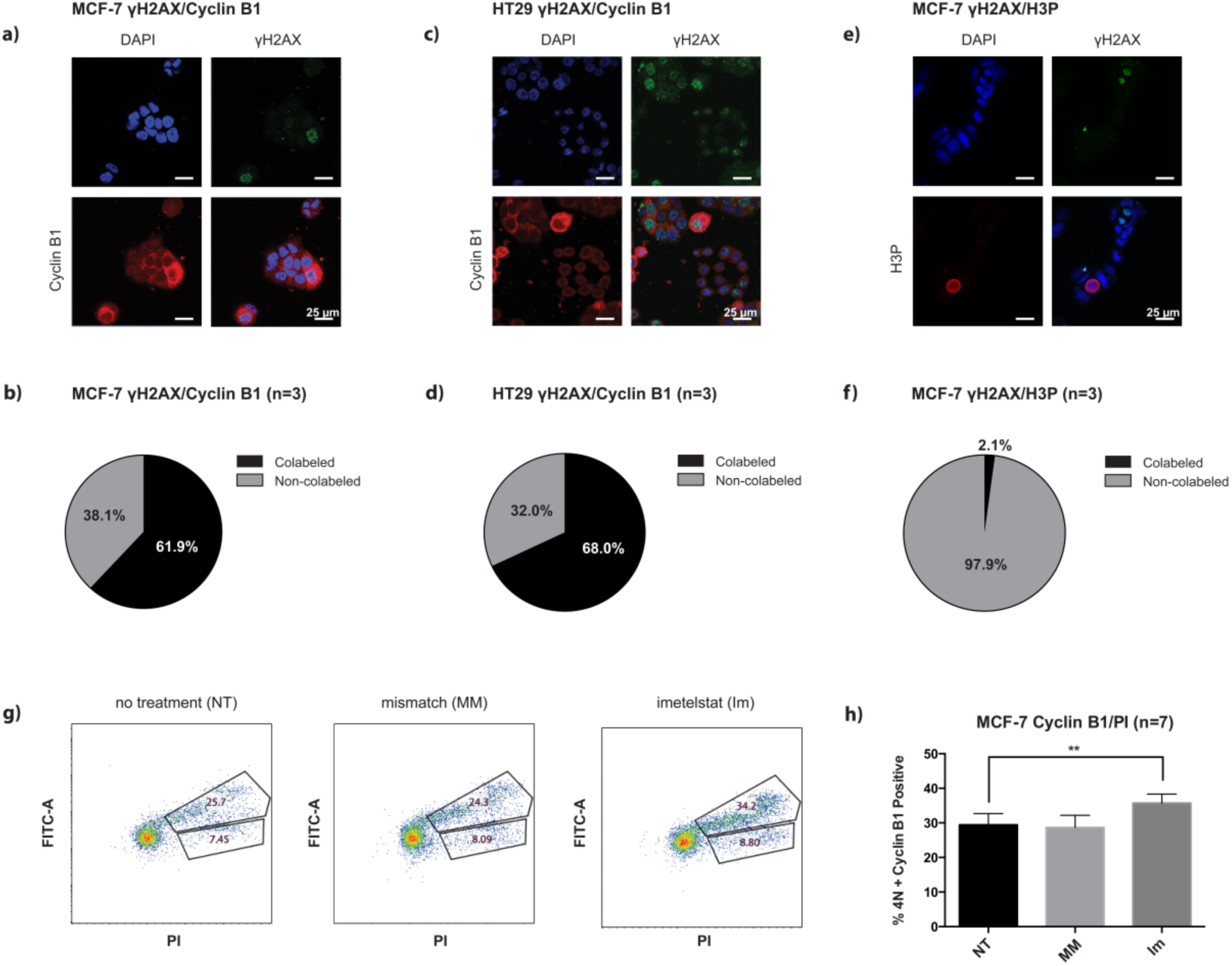
Imetelstat increases the population of cells in late S/G2 phases. (a-c) Following 24 h treatment with imetelstat (10 µM), MCF-7 and HT29 cells were labeled for γH2AX in combination with cytoplasmic cyclin B1, a marker of late S/G2 phases, or phospho-histone H3 (H3P), a marker of M phase. (d-f) DNA damage foci-positive cells (≥10 foci) co-labeled with cyclin B1 or H3P were quantified, with >400 cells analyzed for each condition. (g-h) MCF-7 cells were treated with imetelstat (Im), the mismatch oligo (MM) or no drug (NT) for 24 h then labeled with FITC-conjugated anti-cyclin B1 and propidium iodide (PI). ANOVA and Fisher’s LSD test were used to generate P-values (* = P ≤0.05, ** = P ≤ 0.01). Error bars represent SD.

To confirm the telomeric origin of imetelstat-induced γH2AX DDR foci, we performed chromatin-immunoprecipitation (ChIP) using anti-γH2AX antibody. In agreement with our immunocytochemistry observations, we confirmed increased γH2AX ChIP signals following 24h imetelstat and etoposide treatments, but not after treatments with the mismatch oligo or vehicle control (Fig. 2 g-i). Quantitative PCR measurements of γH2AX ChIP signals showed enrichment of the telomeric sequence following both imetelstat and etoposide treatments (Fig. 2 g). In contrast, γH2AX signal enrichment at Alu-repeat sequences was only observed with etoposide treatments (Fig. 2 h). When γH2AX signal enrichment at telomeres was normalized to the signals measured with the Alu-repeat reference, the imetelstat treatment group was significantly different from the other three treatment groups, suggesting that imetelstat induced the formation of telomere-specific γH2AX loci (Fig. 2 i). This accumulation of γH2AX DDR foci at telomeres following imetelstat treatment is consistent with an increase in uncapped telomeres caused by the absence or inhibition of telomerase activity.

In order to assess the distribution of γH2AX DDR foci-positive cell populations and their corresponding cell cycle phase, using immunocytochemistry, we co-labeled imetelstat-treated MCF-7 and HT29 cells with DDR and cell cycle phase markers. Since γH2AX DDR foci-positive cells co-labeled with cytoplasmic cyclin B1, a marker of late S/G2 phases (Fig. 3 a-d), but not phospho-histone H3 (H3P), an M phase marker in MCF-7 cells (Fig. 3 e-f), the γH2AX DDR foci-positive cells were concluded to be residing in the S/G2 phases.

Consistent with the immunocytochemistry data, FACS measurements showed the proportion of MCF-7 cells that were both cyclin B1-positive and contained 4N DNA content increased with imetelstat treatment, relative to no drug treatment and/or mismatch oligo controls (Fig. 3 g-h). Considering that the magnitude of this increase was small, a delayed clearance from, rather than a complete arrest at, G2 phase was more likely. Overall, our data indicate that imetelstat-induced telomerase inhibition correlates with an accumulation of DDR marker γH2AX and a delayed clearance of the telomere checkpoint at G2 phase of the cell cycle.

### ATM inhibition abolishes the increase in 4N DNA cells caused by imetelstat treatment

As ATM activation is concurrent with the resolution and rebuilding of telomere structures, we next examined whether the effects of imetelstat on cell cycle progression depend upon active ATM signaling. We applied pharmacological inhibitor order-of-addition treatment regimens using imetelstat (Im) and the ATM-specific inhibitor KU55933 (Ku). The treatment conditions tested include: single drug treatments of imetelstat (Im|Im) or KU55933 (Ku|Ku) for 48 h; combinational treatment of imetelstat for 24 h then both imetelstat and KU55933 for another 24 h (Im|Im + Ku); and combinational treatment of KU55933 for 24 h then both imetelstat and KU55933 for another 24 h (Ku|Im + Ku) (Fig. 4 a).

**Figure 4.**
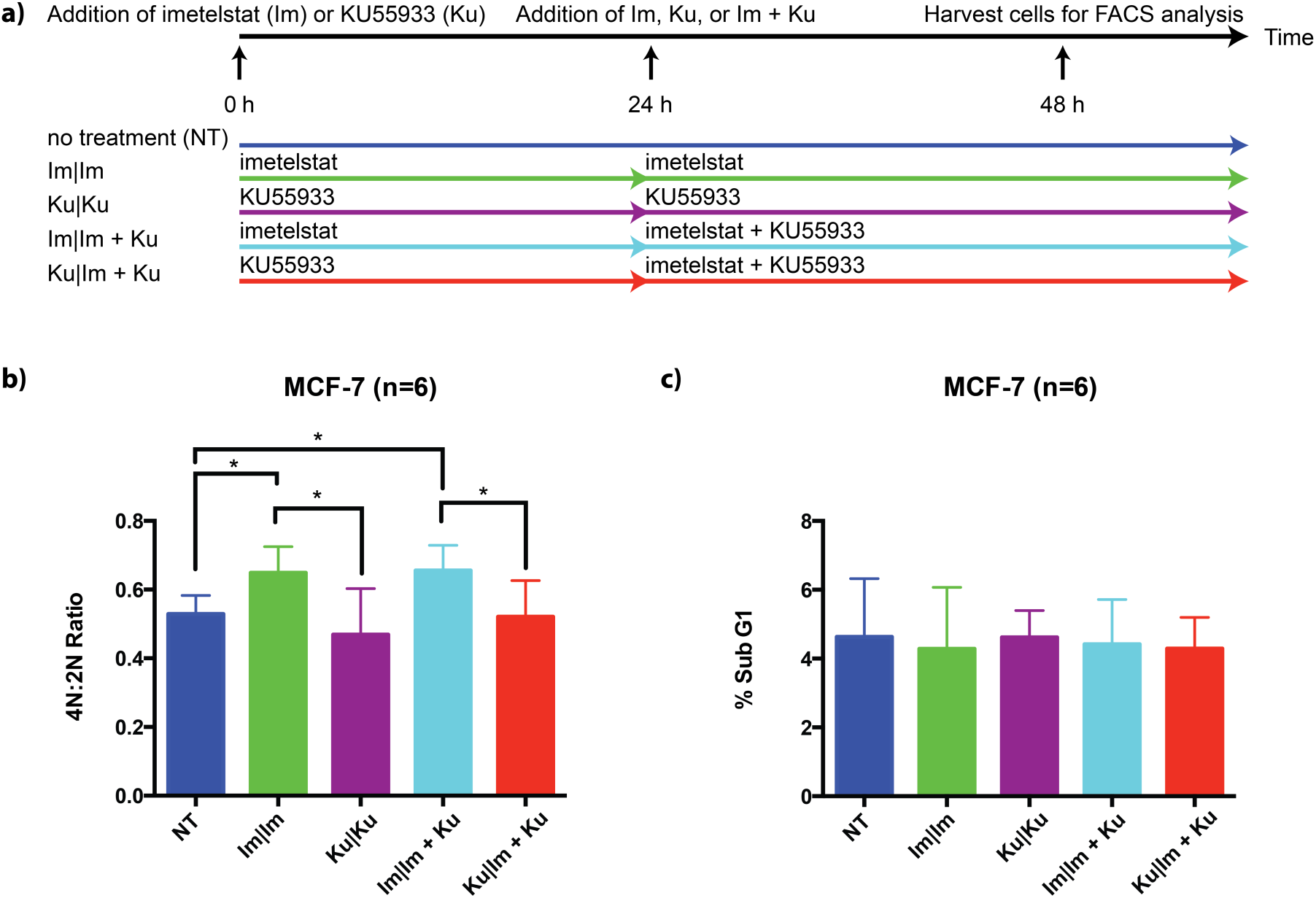
KU55933 and imetelstat affects cell cycle population distribution. (a) MCF-7 cells were treated with the indicated inhibitors (first 24 h | second 24 h) for 48 h. FACS analysis with propidium iodide staining was used to determine the 4N to 2N ratio. (b) Imetelstat (Im|Im) and imetelstat in combination with KU55933 (Im|Im + K) increased 4N DNA cell populations compared to no treatment (NT). Treatment with KU55933 in the first 24 h decreased 4N DNA cell populations compared to the respective treatment regimens using imetelstat in the first 24 h. (c) No difference in sub-G1 populations, a readout of apoptosis and cell death, was observed among cell lines undergoing different treatment regimens. ANOVA and Fisher’s LSD test were used to generate P-values (* = P ≤0.05). Error bars represent SD.

Consistent with Figure 1 a, imetelstat (Im|Im) treatment in MCF-7 cells increased the population of cells with 4N DNA content relative to the population distribution in untreated cells (Fig. 4 b). Notably, this effect on cell cycle progression was abrogated by KU55933 pretreatment (Ku|Im + Ku) but not by treatment with KU55933 after imetelstat exposure (Im|Im + Ku). Treatment with ATM inhibitor alone (Ku|Ku) resulted in a reduction in the cell population with 4N DNA content, but this reduction was not statistically significant in our data analysis.

Treatment-induced cellular toxicities can also affect cell cycle profiles. To differentiate this possibility from phase-specific cell cycle regulation, we analyzed sub-G1 populations by FACS as a functional readout of cell death at the time of DNA content analysis. The proportions of sub-G1 cell populations were not significantly affected by single imetelstat (Im|Im) and KU55933 (Ku|Ku) treatments, or by any treatment combinations, indicating that these changes in cell cycle profiles were unlikely to be caused by increased rates of cell death (Fig. 4 c).

Additionally, we assessed whether KU55933 interfered with the efficiency of imetelstat-induced telomerase inhibition by measuring telomerase activity with TRAP. Cells pre-treated with KU55933 (Ku|Im + Ku) also showed no telomerase activity in the presence of imetelstat (Supplementary Fig. 4), confirming that ATM inhibition did not affect imetelstat’s ability to completely inhibit telomerase actions. Together, these results imply that cell cycle alterations caused by imetelstat-induced telomerase inhibition are dependent on functional ATM signaling.

### Transient formation of telomere dysfunction signals depend upon active ATM signaling

Imetelstat was observed to delay G2 checkpoint clearance in an ATM-activity dependent manner. To connect the role of ATM in G2 checkpoint regulation with imetelstat-dependent γH2AX DDR foci formation, we again used our order-of-addition experimental scheme coupled with immunofluorescence detection of DDR signals. MCF-7 cells were treated with KU55933 and imetelstat for 48 h, either alone or in different combinations, and γH2AX DDR foci were quantified. As in our previous experiments, imetelstat treatment (Im|Im) resulted in an increase in γH2AX DDR foci relative to the untreated (NT) negative control (Fig. 6 a-b). Also as expected, we observed that treatment with KU55933 alone (Ku|Ku) or before imetelstat addition (Ku|Im + Ku) abolished γH2AX DDR foci formation. Interestingly, treatment with KU55933 after 24 h incubation with imetelstat (Im|Im + Ku) only showed a slight, but not statistically significant, reduction in γH2AX DDR foci formation. It is conceivable that blocking the activity of ATM prevents the phosphorylation and propagation of new γH2AX foci but does not remove the phosphorylated proteins that have already formed (Fig. 5 a-b).

**Figure 5.**
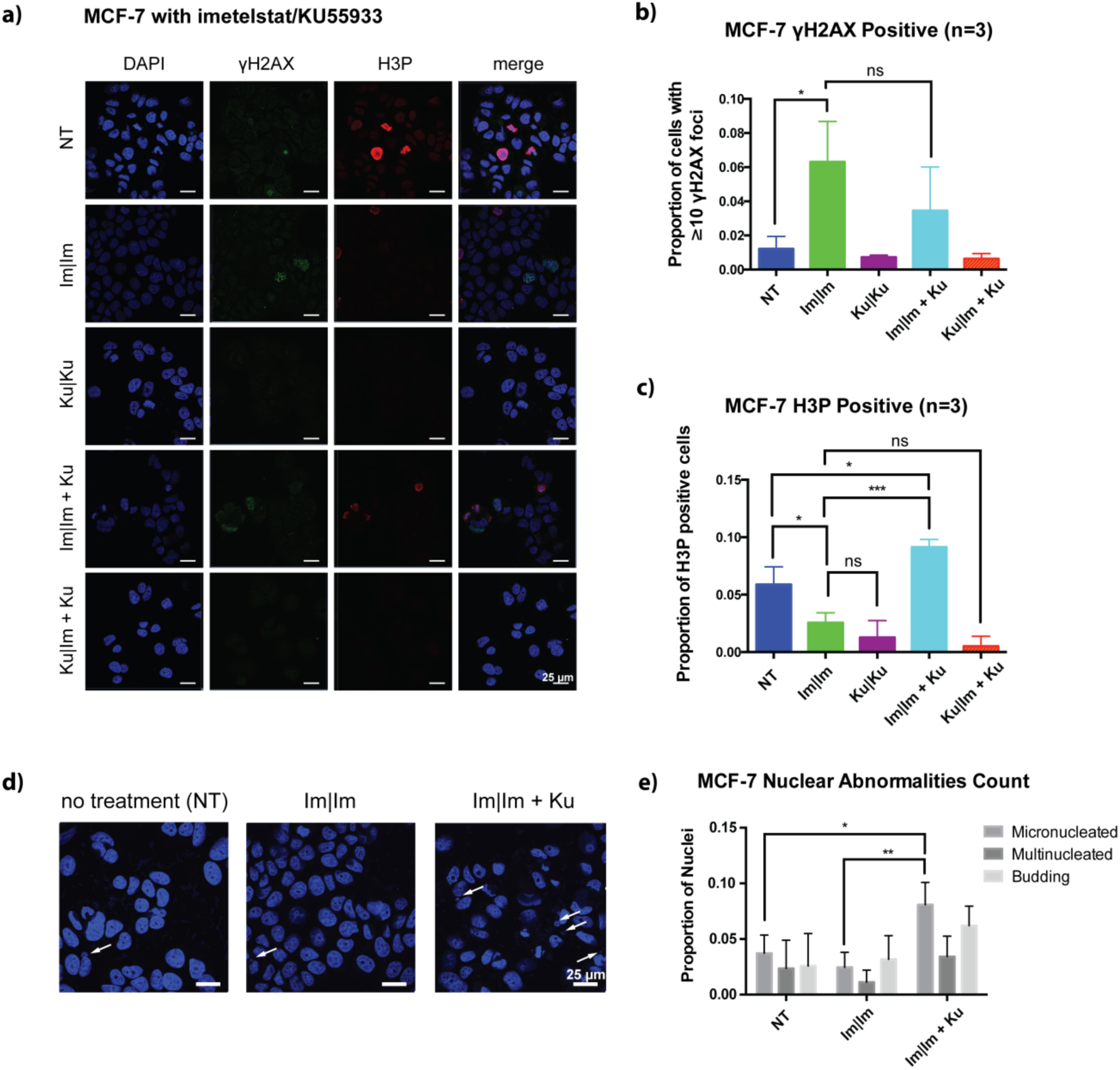
Functional ATM signaling is required for imetelstat-induced DNA damage foci formation and the G2/M checkpoint stall. (a) MCF-7 cells were treated with the indicated inhibitor(s) for 48 h using the same scheme as Fig. 2 before γH2AX and phospho-histone H3 (H3P) labeling. (b-c) Images were scored blindly to quantify co-labeling, with >400 cells scored per condition. ANOVA and Fisher’s LSD test were used to generate P-values (* = P ≤0.05, ** = P ≤ 0.01, *** = P ≤ 0.001). Error bars represent SD. (d) Representative images of DAPI nuclear staining for treatments examined. Arrows indicate micronuclei. (e) Nuclear abnormalities were quantified from the same sets of images as in Fig. 5. Error bars represent SD. ANOVA and Fisher’s LSD test were used to generate P-values (* = P ≤0.05, ** = P ≤ 0.01, *** = P ≤ 0.001, **** = P ≤ 0.0001).

To provide further insight into the fate of imetelstat-treated cells that escape G2-stalling, we labeled cells from different treatment groups for phospho-histone H3P and quantified the proportion of cells progressing to M phase (Fig. 5 a, c). When no drug treatment was administered, ∼5% of MCF-7 cells were H3P-positive. Treatment with imetelstat alone (Im|Im) decreased the proportion of cells in M phase, consistent with a stall at the G2 checkpoint due to persistent DDR signaling. In contrast, cells treated with imetelstat and then KU55933 (Im|Im + Ku) showed an increase in the proportion of cells in M phase, suggesting that continuous ATM signaling is essential for the G2 stall caused by imetelstat-induced DDR signaling. Blocking ATM signaling following transient telomerase inhibition likely released MCF-7 cells from the G2 checkpoint and allowed progression to the next phase of the cell cycle. This release of previously stalled cells manifested as an increase in cells entering M phase. This treatment regimen was also associated with the observation of increased micronuclei formation that indicated mitotic defects and increased genomic instability (Fig. 5 d-e).

Similar to continuous imetelstat treatment, treatment with KU55933 alone (Ku|Ku) or prior to imetelstat addition (Ku|Im + Ku) reduced the number of M-phase cells, suggesting that the order of ATM and telomerase inhibition is important. Consistent with the relative decrease in 4N DNA cell populations observed in our earlier FACS experiments (Fig. 4 b), ATM inhibition may induce faster passage through G2/M phases using multiple parallel mechanisms (see discussion) such that the effects of telomerase inhibition are masked. Our data demonstrate that ATM inhibition before imetelstat treatment removed the effects of telomerase-inhibition–induced G2 stalling, confirming that functional ATM activity is essential for the telomere checkpoint.

### Imetelstat potentiation of etoposide cytotoxicity depends upon active ATM signaling

Previously, we observed that imetelstat potentiated the cytotoxicity of S/G2-specific DNA-damaging agents, including the topoisomerase inhibitors etoposide and irinotecan (5). Addition of ATM inhibitor KU55933 following 24 h of imetelstat treatment (Im|Im + Ku) further increased the cytotoxicity of etoposide. However, the cytotoxic effects from inhibition of ATM signaling before imetelstat treatment (Ku|Im + Ku) were not tested in this previous study. To clarify the role of ATM signaling in imetelstat-induced potentiation of etoposide cytotoxicity, we again performed order-of-addition treatment experiments using the colony forming unit assay (5-7,11,14). MCF-7 cells were pre-treated for 24 h with imetelstat or KU55933 before the addition of etoposide with continued inhibitor treatment or in combination with both imetelstat and KU55933 for 24 h. After 48 h of treatment, cells were harvested and set in soft agar medium to recover for 2 weeks (Fig. 6 a).

**Figure 6.**
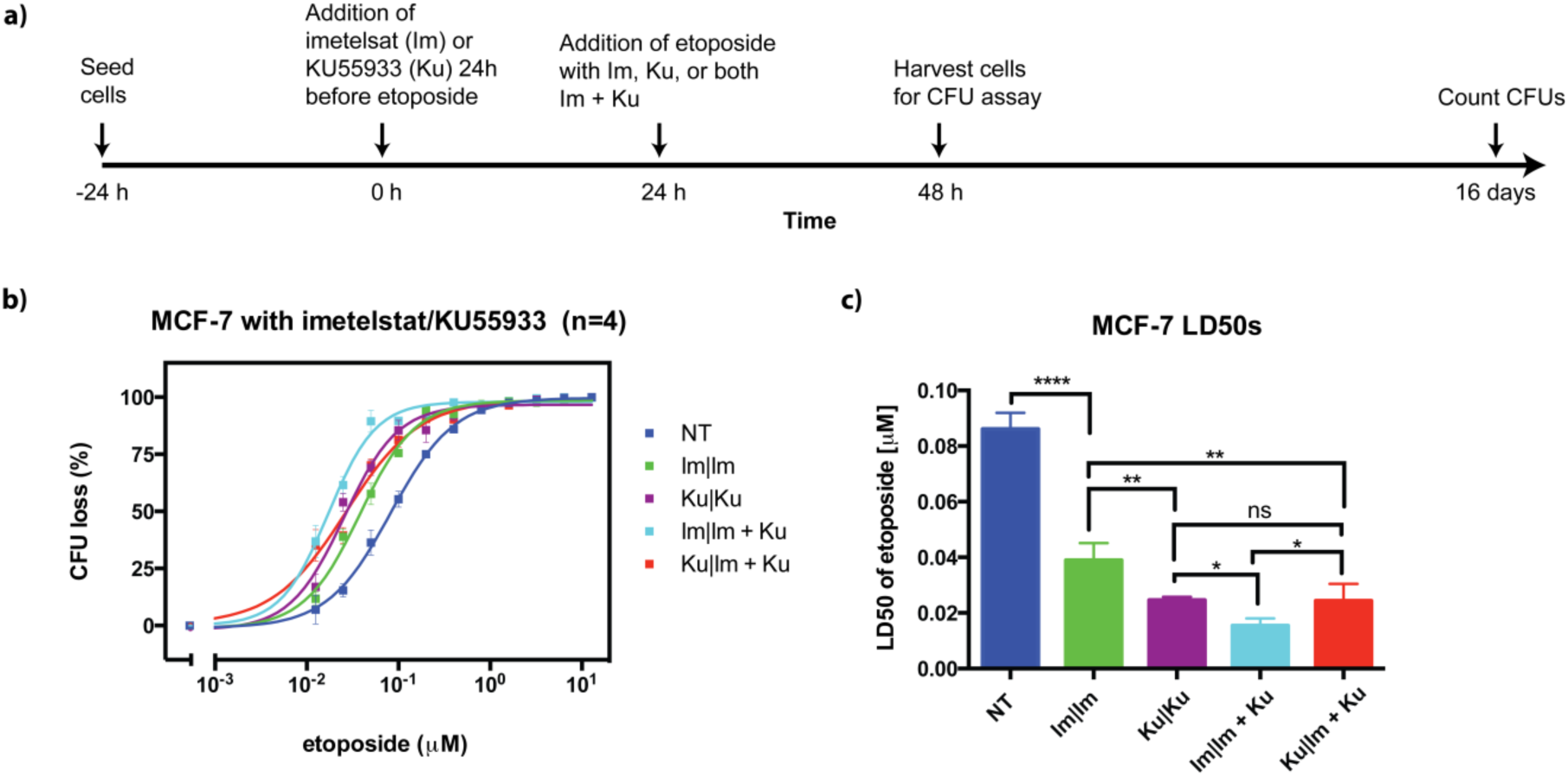
Imetelstat potentiation of etoposide cytotoxicity requires functional ATM signaling. (a) MCF-7 cells were treated with imetelstat (10 µM) or KU55933 (10 µM) for 24 h prior to addition of serial dilutions of etoposide (12.8 – 0.0125 µM) in combination with the previously used inhibitor or in combination with both inhibitors (Im + Ku). After 24 h incubation with etoposide, cells were counted and seeded in soft agar using the colony formation assay (CFU). Colonies were scored after 14 days. (b) Dose-response curves of MCF-7 cells given the indicated inhibitors (first 24 h | second 24 h in combination with etoposide) or no treatment (NT) before etoposide addition. (c) LD_50_s obtained from dose-response curves. Error bars represent SEM.

In agreement with previous results, imetelstat (Im|Im) or KU55933 (Ku|Ku) treatment alone significantly potentiated etoposide cytotoxicity in MCF-7 cells (Fig. 6 b-c) (5,10-11,13,18-19,27). ATM inhibition consistently resulted in greater potentiation of etoposide cytotoxicity than imetelstat-induced telomerase inhibition alone. This observation was attributed to the broader role of ATM in signal transduction regulation of cellular responses to double-strand DNA damage (21), in addition to its role in telomere maintenance and structural homeostasis.

Consistent with our previous data, an additive effect on etoposide cytotoxicity was observed when the cells were treated with both inhibitors in combination following imetelstat pre-treatment (Im|Im + Ku). In contrast, the additive effect was lost when cells were pre-treated with KU55933 before imetelstat and KU55933 addition (Ku|Im +Ku). These observations suggest ATM-dependent DDR signaling may be required for the potentiation of etoposide cytotoxicity by telomerase inhibition.

Notably, the rank order of treatment conditions leading to increased sensitivities towards etoposide mirrors the rank order of treatment conditions leading to increased DDR foci accumulation observed in our previous FACS experiments. The most cytotoxic treatment condition (Im|Im + Ku) was also observed to increase the proportion of cells in M phase in our previous ICC experiments. Based on these observations, we reasoned that the increased cytotoxicity might be linked to increased mitotic errors caused by the premature release of chromosomes with unstructured telomeres from G2 checkpoint. This was confirmed by our previous observation of increased micronuclei formation with the treatment condition (Im|Im + Ku) indicating mitotic defects and increased genomic instability (Fig. 5 d-e). This increase in the number of cells with mitotic defects acts as an additional cytotoxic mechanism in etoposide-treated MCF-7 cells.

## DISCUSSION

### TERT addiction manifests as a kinetic advantage in cell cycle progression

The role of telomerase in telomeric chromatin formation is inferred through its *de novo* telomere synthesis function but has never been directly demonstrated. Our data suggest that telomerase improves the resolution of transient telomere dysfunction signals, thereby facilitating efficient higher-order telomeric chromatin formation and clearance from the G2 checkpoint.

Telomere chromatin formation requires the creation of telomeric G-rich overhangs that invade the double-stranded telomeric DNA region and form displacement loops (1,13). In primary cells, telomerase is inactive and the generation of G-rich overhangs and T-loops is dependent upon C-strand-specific nuclease activity (1,11,13-16,28). In cancer cells, telomerase expression provides another avenue for the de novo synthesis of G-rich telomeric DNA repeats, in addition to nuclease activity (1,11,13-16,28). This may explain our observation that telomerase inhibition delayed, but did not stop, cell cycle progression. In telomerase-positive cells, imetelstat treatment blocked the access of reverse transcriptase to its telomeric DNA substrate, thereby preventing *de novo* synthesis of G-rich telomere repeats (11,13-17,28-29). Consequently, telomere dysfunction signals at unstructured telomeres took longer to resolve and resulted in delayed passage through cell cycle checkpoints and a temporary accumulation of cells in G2 phase, although the lack of telomerase action did not entirely arrest cells at G2 phase. This may be because G-rich overhangs for T-loop formation could still be produced, albeit with reduced efficiency, through the actions of multiple nucleases such as Exo1 (10-12,14,30). However, nuclease recruitment and processing may have slower kinetics than telomere repeat synthesis, as reflected by the delay in cell cycle progression (13,28). From this, we inferred that telomerase catalysis is more efficient than the actions of C-strand-specific nucleases at G-rich overhang formation.

The observation that telomerase improved the kinetics of telomere dysfunction signal clearance explains the selective growth advantage of telomerase-positive cells. The faster cell cycle progression may cause cancer cells to become “addicted” to telomerase activity and explain the preponderance of telomerase overexpression (>85%) for telomere maintenance over the alternate lengthening of telomere mechanism in surveyed cancers (26). As telomerase-negative cells can reform T-loops using C-strand-specific nucleases, their cell cycle progression is predicted to be unaltered by imetelstat treatment (5,7,8,20,23-24,28). This prediction is supported by the unchanged cell cycle profiles (Fig. 1) and the lack of significant decreases in cell growth in imetelstat-treated telomerase-negative cell lines (Supplementary Fig. 1 c). Therefore, somatic tissues with low telomerase expression are also predicted to have tolerable clinical toxicity profiles for telomerase inhibition by imetelstat (1,5-7,17).

### ATM signaling is necessary for telomerase actions at unstructured telomere ends

Functional ATM is required for normal elongation of telomeres by telomerase in human cells (5,10-11,13,18-19,31-33). We observed that ATM inhibition prior to imetelstat treatment abolished the delay in cell cycle progression induced by telomerase inhibition. This observation may be due to the inhibition of telomerase recruitment to the telomere in the absence of ATM activity (34) or the inhibition of a telomerase-independent ATM function that leads to changes in G2 checkpoint engagement (35).

Notably, ATM inhibition was previously shown to accelerate passage through G2 phase by the removal of inhibitory phosphorylation of C-strand-specific Exo1 nuclease (29). Increased Exo1 activity leads to faster G-rich overhang formation and thus, increases the kinetics of T-loop formation in the absence of telomerase actions. Accordingly, accelerated G2/M progression is consistent with the observed trend of reduced populations of cells with 4N DNA content following KU55933 treatment (Fig. 4 b).

Interestingly, reversal of the order of inhibitor addition, with the addition of imetelstat before ATM inhibition, resulted in increases in M-phase cell populations. This suggests that the ATM-dependent telomere checkpoint was quickly engaged following telomerase inhibition. Subsequent ATM inhibition released previously stalled cells from G2 phase to M phase, confirming that continuous ATM signaling is crucial for the maintenance of this checkpoint (36). In M phase, uncapped chromosome ends resulted in abnormal cell division and/or mitotic catastrophe (Fig. 6 d-e). The higher levels of micronuclei formation indicate possible mitotic defects and chromosomal instability due to errors in telomere processing.

### Therapeutic implications of short-term telomerase inhibition

Previous work in our laboratory demonstrated that combining telomerase inhibition by imetelstat with genotoxic agents potentiated the cytotoxic effects of G2-specific DNA-damaging agents such as topoisomerase inhibitors (5,10-11). Data from our current study indicate that this potentiating effect is likely due to an ATM-dependent DNA-damage signal induced by telomerase inhibition. The telomere dysfunctions signals at unstructured chromosome ends may act in an additive manner with G2-specific DNA-damaging agents such that a greater proportion of cells pass the apoptotic threshold. Our hypothesis is supported by previous observations that telomerase depletion in yeast caused chronic replication stress and stalled replication forks (30). In this context, topoisomerase inhibitors and other drugs that cause replicative stress may synergize particularly well with imetelstat.

The potentiation of topoisomerase inhibitors by imetelstat may also be due to pharmacodynamic interactions between telomerase inhibition and topoisomerase inhibition. As previously observed, proper T-loop formation may involve topoisomerase activity (7). The loss of telomerase activity, and thus delay in clearance of telomere dysfunction signals and T-loop formation, in combination with the absence of topology relief may further damage telomere structures and result in cytotoxicity responses. This may provide another mechanism by which imetelstat can potentiate topoisomerase inhibitor cytotoxicity.

In order for telomeres to shorten significantly, multiple rounds of cellular replication are required. The long lag time associated with telomere shortening has been a major theoretical barrier to the utilization of telomerase inhibitors for anti-cancer chemotherapy (1). Recent clinical trials of imetelstat in myelofibrosis and thrombocythemia have cast doubt on this premise. Telomere length did not change in response to therapy and baseline telomere length was not predictive of a positive therapeutic response (8,31,37). In this context, our data provide an alternate explanation for the observed clinical effects of imetelstat: telomerase inhibition in these hematopoietic cell types may induce distortions in cell cycle kinetics without parallel observable effects on telomere-length regulation.

Telomeric-DNA-replication stress due to telomerase inhibition may be partially relieved by an increase in the dNTP (purine) pool, as suggested in previous studies (6,7,30). However, hematopoietic cancers frequently display dysregulated dNTP metabolism (10,13,18-19,38). Therefore, this model predicts that existing therapeutic agents targeting the available dNTP pools, such as mycophenolic acids, will have synergistic effects with imetelstat treatments in vulnerable hematopoietic cell types.

We observed that transient telomerase inhibition by imetelstat delayed the clearance of ATM-dependent telomere dysfunction signals that engage the G2 checkpoint, resulting in altered cell cycle kinetics. Through the delayed resolution of telomere dysfunction signals, imetelstat also sensitized telomerase-positive cells to G2-specific DNA-damaging agents. These observations allude to a separate mechanism by which telomerase inhibition could affect telomere-maintenance kinetics and homeostasis. Our data is relevant for understanding the role of telomerase in the formation of higher-order telomeric-chromatin structures and cell cycle progression, thereby presenting new testable hypotheses and possibilities for combination drug regimens.

## MATERIALS AND METHODS

### Cell lines and reagents

MCF-7, MDA-MB 231, HT29, LS180, BJ fibroblasts, and WI-38 VA-13 were obtained from the American Type Culture Collection (ATCC). MDA-MB 231 (NucLight Red) cells were obtained from Essen Bioscience. Cell culture media, antibiotics, and other cell culture reagents were purchased from Invitrogen/Life Technologies unless otherwise noted. Cells were maintained under standard culture conditions of 37°C and 5% CO_2_ with penicillin and streptomycin antibiotics (100 U each) and in the presence of appropriate fetal bovine serum (FBS) concentrations (5-15%), as indicated by the ATCC.

Etoposide (Sigma/Aldrich) was administered in dose-response treatments in 2-fold serial dilution and a maximum dose of 10uM. ATM inhibitor KU55933 (Calbiochem) was administered as 10 µM in DMSO, a concentration previously determined to efficiently inhibit ATM function (5,20,23). Imetelstat and its mismatch oligo control (MM) were obtained from Geron, resuspended in PBS, and stored at -20°C. Stock concentrations were determined before each experiment using UV-spectrophotometer absorbance. Imetelstat and MM were administered at 10 µM, a dose previously determined to inhibit telomerase activity by >100 fold in multiple cancer cell types (5).

### Colony forming unit assay (CFU)

The assay was performed as previously described (1,5). Cells were seeded and allowed to grow into individual colonies at 37°C under 5% CO_2_ for 2 weeks. Colonies were counted as positive if colony sizes exceeded 50 µm. Dose-response analysis was performed using GraphPad Prism (v6.0b).

### Immunocytochemistry (ICC)

Labeling was conducted as previously described (5,31-32). Primary antibodies were sourced and diluted as follows: anti-phospho-histone H2AX (Ser139) 1:500 (JBW301 EMD Millipore), phospho-H3 1:500 (06-570 EMD Millipore), and cyclin B1 1:100 (Santa Cruz H-433). Images were collected using a Zeiss LSM 700 confocal microscope and Zen 2012 (Zeiss) software. To quantify DNA-damage response foci, the collected images were blinded and scored with >400 cells analyzed for each condition. Nuclear abnormality scoring was conducted as previously described (39).

### Fluorescence-activated cell sorting (FACS)

Cells were plated and allowed to settle before treatment for 24 to 48 h with the described drug regimens. The treated cells were harvested by trypsinization and fixed with ethanol. FITC-Anti-Cyclin B1 (BD Biosciences) immunolabeling and propidium iodide nucleic acid labeling were conducted according to the manufacturer’s protocol (Becton Dickinson). RNAse treatment was used to remove non-specific signals. Labeled cells were sorted using a BD LSRII flow cytometer (UBC Life Sciences Institute). For each cell sample, a minimum of 10,000 events were recorded. Flow Jo (Tree Star Inc) software and standard gating procedures were used to quantify the sub-G1 population and the number of cells with 2N and 4N DNA content.

### Chromatin-immunoprecipitation (ChIP)

Cells were plated in 10 cm plates and allowed to settle before treatment for 24 h with the described drug regimens. The treated cells were PBS-washed and cross-linked in 1% formaldehyde. Cross-linked samples were collected by scraping in ChIP lysis buffer (20 mM Tris-HCl pH 8.0, 140 mM NaCl, 1 mM EDTA pH 8.0, 1% Triton X-100, 0.1% SDS, 1X protease inhibitor cocktail) and sonicated with a Covaris m220 ultrasonicator. The sonicated fractions were then precleared with Protein G sepharose beads (GE Healthcare, #17-0618) and then subjected to immunoprecipitation (IP) with 3 µg of anti-phospho-Histone H2A.X (Millipore 05-636). IP with 30 µL of BSA-preblocked protein G beads (without primary antibody) was used as control. Chromatin immunoprecipitates were washed sequentially in ChIP Wash Buffer A (0.1% SDS, 1% Triton X-100, 2 mM EDTA, 20 mM Tris-HCl pH 8.0, 150 mM NaCl), ChIP Wash Buffer B (0.1% SDS, 1% Triton X-100, 2 mM EDTA, 20 mM Tris-HCl pH 8.0, 500 mM NaCl), ChIP Wash Buffer C (0.25 M LiCl, 1% NP-40, 1% Sodium Deoxycholate, 1 mM EDTA, 10 mM Tris-HCl pH 8.0), and lastly TE buffer. The ChIP samples were reversed cross-linked overnight in 65°C, and extracted DNA was purified with a DNA cleanup kit (BioBasic). Quantitative PCR was performed using a Tel1b (telomere sequence) primer set (Forward: CGGTTTGTTTGGGTTTGGGTTTGGGTTTGGGTTTGGGTT, Reverse: GGCTTGCCTTACCCTTACCCTTACCCTTACCCTTACCCT) and an Alu repeat reference primer set (Forward: GACCATCCCGGCTAAAACG, Reverse: CGGGTTCACGCCATTCTC) (40).

### Data analysis

GraphPad Prism version 6 (GraphPad Software Inc, San Diego, CA) was used for statistical analysis and data presentation. Student’s *t*-tests or, where appropriate, ANOVA followed by Fisher’s LSD test were used to generate P-values. *P* < 0.05 was considered statistically significant.

## ACKNOWLEDGEMENTS

We thank Geron Corporation for the gift of imetelstat. We appreciate assistance from Arthur Chen for quantifying the ICC images, Andy Johnson for FACS use and analysis, and JiaLin Xu for technical help with TRAP experiments. Anna Krassowska, Neeru Batra from Geron, and members of the Wong Lab are thanked for critically reading the manuscript.

## REFERENCES

1. Shay JW. Role of Telomeres and Telomerase in Aging and Cancer. Cancer Discov. 2016 Jun;6(6):584–93.

2. Hukezalie KR, Wong JM. Structure-Function Relationship and Biogenesis Regulation of the Human Telomerase Holoenzyme. FEBS J. 2013 Apr 3.

3. Shay JW, Wright WE. Hayflick, his limit, and cellular ageing. Nat. Rev. Mol. Cell Biol. 2000 Oct;1(1):72–6.

4. Shay JW, Wright WE. Role of telomeres and telomerase in cancer. Semin. Cancer Biol. 2011 Dec;21(6):349–53.

5. Tamakawa RA, Fleisig HB, Wong JMY. Telomerase inhibition potentiates the effects of genotoxic agents in breast and colorectal cancer cells in a cell cycle-specific manner. Cancer Res. 2010 Nov 1;70(21):8684–94.

6. Martínez P, Blasco MA. Telomeric and extra-telomeric roles for telomerase and the telomere-binding proteins. Nat. Rev. Cancer. 2011 Mar;11(3):161–76.

7. Giraud-Panis M-J, Pisano S, Benarroch-Popivker D, Pei B, Le Du M-H, Gilson E. One identity or more for telomeres? Front Oncol. 2013;3:48.

8. Wong JMY, Collins K. Telomere maintenance and disease. Lancet. 2003 Sep 20;362(9388):983–8.

9. Shay JW, Bacchetti S. A survey of telomerase activity in human cancer. Eur. J. Cancer. 1997 Apr;33(5):787–91.

10. García-Cao M, O’Sullivan R, Peters AHFM, Jenuwein T, Blasco MA. Epigenetic regulation of telomere length in mammalian cells by the Suv39h1 and Suv39h2 histone methyltransferases. Nat. Genet. 2004 Jan;36(1):94–9.

11. Arnoult N, Karlseder J. Complex interactions between the DNA-damage response and mammalian telomeres. Nat. Struct. Mol. Biol. 2015 Nov;22(11):859–66.

12. Kibe T, Zimmermann M, de Lange T. TPP1 Blocks an ATR-Mediated Resection Mechanism at Telomeres. Mol. Cell. 2016 Jan 21;61(2):236–46.

13. Lenain C, Bauwens S, Amiard S, Brunori M, Giraud-Panis M-J, Gilson E. The Apollo 5′ exonuclease functions together with TRF2 to protect telomeres from DNA repair. Curr. Biol. 2006 Jul 11;16(13):1303–10.

14. Martínez P, Blasco MA. Replicating through telomeres: a means to an end. Trends Biochem. Sci. 2015 Sep;40(9):504–15.

15. Zhong FL, Batista LFZ, Freund A, Pech MF, Venteicher AS, Artandi SE. TPP1 OB-fold domain controls telomere maintenance by recruiting telomerase to chromosome ends. Cell. 2012 Aug 3;150(3):481–94.

16. Nandakumar J, Cech TR. Finding the end: recruitment of telomerase to telomeres. Nat. Rev. Mol. Cell Biol. 2013 Feb;14(2):69–82.

17. Herbert B-S, Gellert GC, Hochreiter A, Pongracz K, Wright WE, Zielinska D, et al. Lipid modification of GRN163, an N3“-->P5” thio-phosphoramidate oligonucleotide, enhances the potency of telomerase inhibition. Oncogene. 2005 Aug 4;24(33):5262–8.

18. Collins K. The biogenesis and regulation of telomerase holoenzymes. Nat. Rev. Mol. Cell Biol. 2006 Jul;7(7):484–94.

19. Chan SS, Chang S. Defending the end zone: studying the players involved in protecting chromosome ends. FEBS Lett. 2010 Sep 10;584(17):3773–8.

20. Verdun RE, Karlseder J. The DNA damage machinery and homologous recombination pathway act consecutively to protect human telomeres. Cell. 2006 Nov 17;127(4):709–20.

21. Shiloh Y, Ziv Y. The ATM protein kinase: regulating the cellular response to genotoxic stress, and more. Nat. Rev. Mol. Cell Biol. 2013 Apr;14(4):197–210.

22. Burgess DJ, Doles J, Zender L, Xue W, Ma B, McCombie WR, et al. Topoisomerase levels determine chemotherapy response in vitro and in vivo. Proc. Natl. Acad. Sci. U.S.A. 2008 Jul 1;105(26):9053–8.

23. Verdun RE, Crabbe L, Haggblom C, Karlseder J. Functional human telomeres are recognized as DNA damage in G2 of the cell cycle. Mol. Cell. 2005 Nov 23;20(4):551–61.

24. Takai H, Smogorzewska A, de Lange T. DNA damage foci at dysfunctional telomeres. Curr. Biol. 2003 Sep 2;13(17):1549–56.

25. Asai A, Oshima Y, Yamamoto Y, Uochi TA, Kusaka H, Akinaga S, Yamashita Y, Pongracz K, Pruzan R, Wunder E, Piatyszek M, Li S, Chin AC, Harley CB, Gryaznov S. A novel telomerase template antagonist (GRN163) as a potential anticancer agent. Cancer Res 2003 Jul 15;63(14):3931–9.

26. Bryan TM, Englezou A, Dalla-Pozza L, Dunham MA, Reddel RR. Evidence for an alternative mechanism for maintaining telomere length in human tumors and tumor-derived cell lines. Nat. Med. 1997 Nov;3(11):1271–4.),

27. Fleisig HB, Wong JMY. Telomerase promotes efficient cell cycle kinetics and confers growth advantage to telomerase-negative transformed human cells. Oncogene. 2012 Feb 23;31(8):954–65.

28. Wu P, Takai H, de Lange T. Telomeric 3′ overhangs derive from resection by Exo1 and Apollo and fill-in by POT1b-associated CST. Cell. 2012 Jul 6;150(1):39–52.

29. Bolderson E, Tomimatsu N, Richard DJ, Boucher D, Kumar R, Pandita TK, et al. Phosphorylation of Exo1 modulates homologous recombination repair of DNA double-strand breaks. Nucleic Acids Res. 2010 Apr;38(6):1821–31.

30. Xie Z, Jay KA, Smith DL, Zhang Y, Liu Z, Zheng J, et al. Early telomerase inactivation accelerates aging independently of telomere length. Cell. 2015 Feb 26;160(5):928–39.

31. Tefferi A, Lasho TL, Begna KH, Patnaik MM, Zblewski DL, Finke CM, et al. A Pilot Study of the Telomerase Inhibitor Imetelstat for Myelofibrosis. N. Engl. J. Med. 2015 Sep 3;373(10):908–19.

32. Jafri MA, Ansari SA, Alqahtani MH, Shay JW. Roles of telomeres and telomerase in cancer, and advances in telomerase-targeted therapies. Genome Med. 2016;8(1):69.

33. Lee SS, Bohrson C, Pike AM, Wheelan SJ, Greider CW. ATM Kinase Is Required for Telomere Elongation in Mouse and Human Cells. Cell Rep. 2015 Nov 24;13(8):1623–32.

34. Tong AS, Stern JL, Sfeir A, Kartawinata M, de Lange T, Zhu X-D, et al. ATM and ATR Signaling Regulate the Recruitment of Human Telomerase to Telomeres. Cell Rep. 2015 Nov 24;13(8):1633–46.

35. Blasina A, Price BD, Turenne GA, McGowan CH. Caffeine inhibits the checkpoint kinase ATM. Curr. Biol. 1999 Oct 7;9(19):1135–8.

36. Shibata A, Barton O, Noon AT, Dahm K, Deckbar D, Goodarzi AA, et al. Role of ATM and the damage response mediator proteins 53BP1 and MDC1 in the maintenance of G(2)/M checkpoint arrest. Mol. Cell. Biol. 2010 Jul;30(13):3371– 83.

37. Baerlocher GM, Oppliger Leibundgut E, Ottmann OG, Spitzer G, Odenike O, McDevitt MA, et al. Telomerase Inhibitor Imetelstat in Patients with Essential Thrombocythemia. N. Engl. J. Med. 2015 Sep 3;373(10):920–8.

38. Kohnken R, Kodigepalli KM, Wu L. Regulation of deoxynucleotide metabolism in cancer: novel mechanisms and therapeutic implications. Mol. Cancer. 2015;14:176.

39. Fleisig HB, Hukezalie KR, Thompson CAH, Au-Yeung TTT, Ludlow AT, Zhao CR, et al. Telomerase reverse transcriptase expression protects transformed human cells against DNA-damaging agents, and increases tolerance to chromosomal instability. Oncogene. 2016 Jan 14;35(2):218–27.

40. Keefe D, Wang F, Robinson LG, Pan XV, Weissman SM, Liu L, Kalmbach KH. Measurement of telomere length at the single cell level. Protoc. Exch. 2017 Jan 5.

